# Catecholaminergic Manipulation Alters Dynamic Network Topology Across Behavioral States

**DOI:** 10.1101/169102

**Authors:** James M. Shine, Ruud L. van den Brink, Dennis Hernaus, Sander Nieuwenhuis, Russell A. Poldrack

## Abstract

The human brain is able to flexibly adapt its information processing capacity to meet a variety of cognitive challenges. Recent evidence suggests that this flexibility is reflected in the dynamic reorganization of the functional connectome. The ascending catecholaminergic arousal systems of the brain are a plausible candidate mechanism for driving alterations in network architecture, enabling efficient deployment of cognitive resources when the environment demands them. We tested this hypothesis by analyzing both task-free and task-based fMRI data following the administration of atomoxetine, a noradrenaline reuptake inhibitor, compared to placebo, in two separate human fMRI studies. Our results demonstrate that the manipulation of central catecholamine levels leads to a reorganization of the functional connectome in a manner that is sensitive to ongoing cognitive demands.

---

A fundamental question facing modern neuroscience is how local computations are integrated across the brain to support the vast repertoire of mammalian behavior and cognition. Convergent results from multi-modal neuroimaging studies^1–4^ have demonstrated that brain activity during cognitive tasks reflects a balance between regional segregation and network-level integration, in which communication across distributed circuits enables fast and effective cognitive performance^1^.

There is growing evidence that ascending catecholaminergic neuromodulatory projections from the brainstem mediate this integration^1,5^. Projections from arousal-related nuclei, such as the noradrenergic locus coeruleus^6^, arborize widely in target regions and putatively alter network architecture by modulating the impact of incoming neuronal input in an activity-dependent manner^7^. Previous neuroimaging studies in humans have highlighted a close relationship between noradrenaline, network topology and cognitive performance^1,8^. Specifically, increased free noradrenaline has been shown to increase the phasic-to-tonic ratio of neuronal firing in both the locus coeruleus and the cortex. As such, neurons that are less tonically active during the un-stimulated state may also simultaneously demonstrate a heightened responsivity to relevant stimuli^9,10^. However, it is not yet known whether manipulating noradrenaline shapes network topology, or indeed whether the effects of noradrenergic function on network topology differ across behavioral contexts.

To test the hypothesis that ascending catecholamines modulate global network topology as a function of behavioral state, we analyzed two separate fMRI datasets in which individuals were scanned following administration of either atomoxetine (ATX), a noradrenergic reuptake inhibitor^11^, or a pharmacologically-inactive placebo. In the first study, subjects were scanned in the task-free ‘resting’ state^12^; in the second, subjects were scanned while performing a cognitively-challenging N-back task^13^. Based on the opposing effects of ATX on functional connectivity observed in these two studies^12,13^, animal studies that highlight differential effects of ATX on phasic versus tonic locus coeruleus activity^9^ and the hypothesized link between noradrenaline and network topology^1,8^, we expected that ATX administration would manifest distinct topological effects as a function of behavioral state.

## Results

### The effect of atomoxetine on the topological signature of the resting state

In the double-blind, placebo-controlled crossover resting-state study^12^, 24 healthy subjects (age = 19-26) underwent fMRI scanning prior to (t = −20 minutes) and following (t = +90 minutes) the administration of either 40mg of ATX or placebo. To estimate time-resolved network topology, we submitted pre-processed BOLD fMRI data from each subject to a pre-registered analysis pipeline that calculates sliding-window connectivity between regional time-series^14^ and then estimates the resulting topological signature of each windowed graph^1^. Specifically, we used a weighted- and signed-version of the Louvain algorithm^15^ to identify tightly connected communities of regions within each temporal window. We then determined how strongly each region was connected to other regions within its own module (quantified using the within-module degree Z-score: W_*T*_) as well as to regions outside of its own module (quantified using the participation coefficient: B_*T*_) over time. The resultant topology can be summarized at the regional level (e.g. to determine which regions were the most integrated during a particular behavioral state), or at the global level using a joint histogram of W_*T*_ and B_*T*_ values (known as a “cartographic profile”). Rightward fluctuations in the density of the cartographic profile along the horizontal (i.e. B_*T*_) axis reflect a more highly integrated functional connectome, and have been shown to relate positively to individual differences in effective cognitive performance^1,16^.

As predicted (https://osf.io/utqq2), the administration of ATX compared to placebo at rest led to a significant reconfiguration of network-level topology (Figure 1a). Specifically, ATX administration caused a global shift towards segregation that was maximal in lateral frontal, frontopolar and occipital cortices, along with the bilateral amygdala (Figure 1b). Increases in free synaptic noradrenaline are known to down-regulate tonic activity within the locus coeruleus, which has a dense expression of inhibitory *α*2-autoreceptors^6,9^. As such, our results might suggest that network topology in the resting state became segregated due to a reduction in the tonic firing rate of the locus coeruleus^9^, an interpretation that is consistent with recent computational models that demonstrate a strong link between the tonic firing rate of the locus coeruleus and functional signatures of brain network activity^17^.

**Figure 1.**
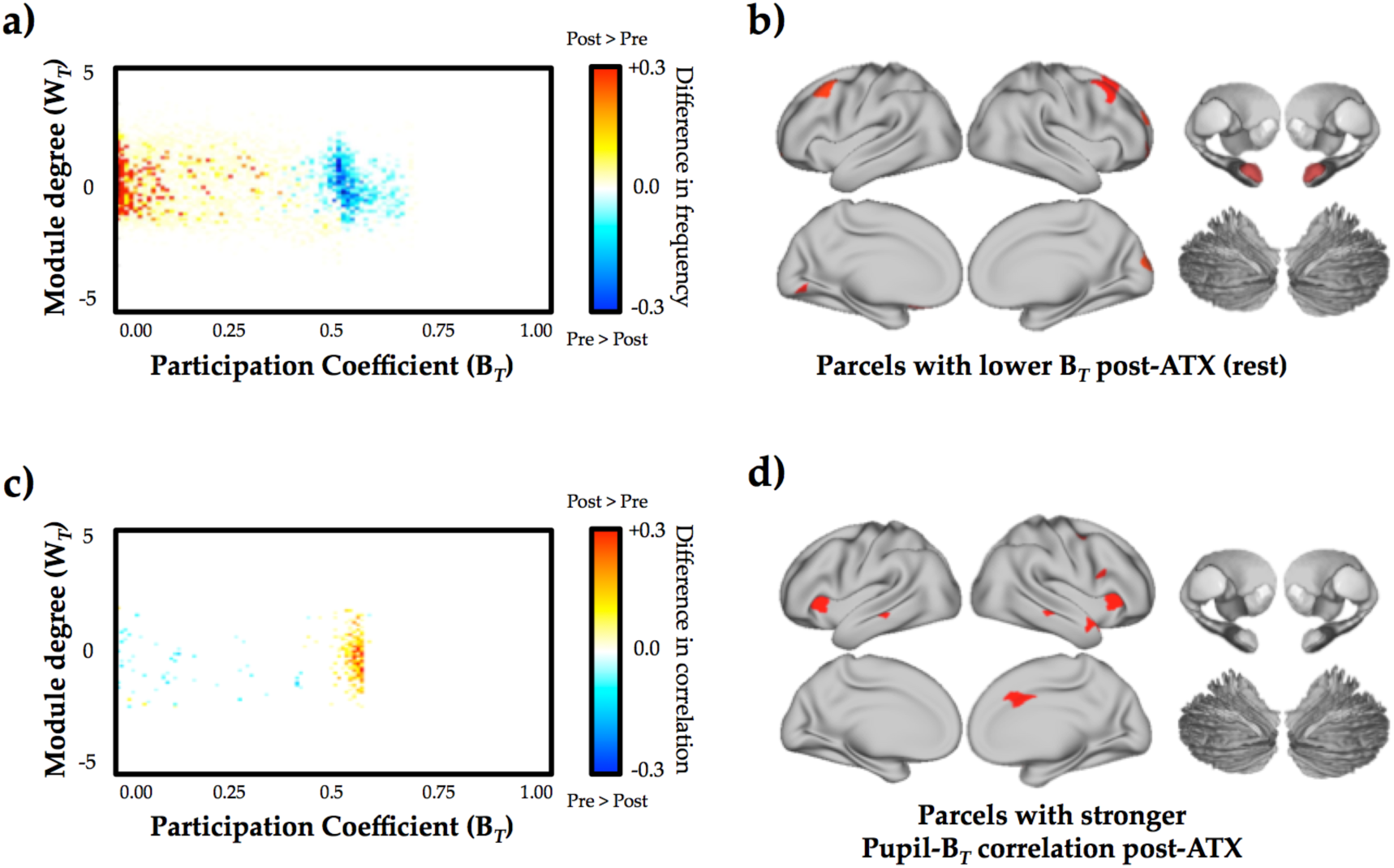
– a) effect of atomoxetine *versus* placebo on the cartographic profile, which demonstrates a shift towards segregation: red/yellow – increased frequency post-atomoxetine and blue – decreased frequency post-atomoxetine (FDR q ≤ 0.05); b) parcels with decreases in their between-module connectivity (i.e. participation coefficient) following atomoxetine (*vs*. placebo) – see Table S1 for parcel MNI co-ordinates (FDR q ≤ 0.05); c) effect of atomoxetine *versus* placebo on the relationship between the cartographic profile and pupil diameter, which demonstrates a shift toward integration: red/yellow – increased frequency post-atomoxetine and blue – decreased frequency post-atomoxetine (FDR q ≤ 0.05); d) parcels with increased time-varying connectivity between between-module connectivity (i.e. participation coefficient) and pupil diameter following atomoxetine (*vs*. placebo) – see Table S1 for parcel MNI co-ordinates (FDR q ≤ 0.05). Key: ATX – atomoxetine; B_*T*_ – between-module connectivity; W_*T*_ – within-module connectivity; see Table S1 for parcel co-ordinates.

### Network topology is sensitive to catecholaminergic manipulation

Although the topological signature observed in the resting state is consistent with a decrease in tonic noradrenaline, *in vivo* experiments in rodents have demonstrated that ATX administration also enhances phasic firing patterns in the locus coeruleus^9^. This in turn should be expected to potentiate phasic noradrenergic responses and hence, integrate the brain, however only under conditions necessary to elicit phasic noradrenergic signaling, such as sensory salience^18^ and acute stress^19^. Thus, in the context of an increase in free noradrenaline, we might expect that the strength of the relationship between network topology and phasic noradrenaline should increase following ATX administration. That is, the presence of extra noradrenaline should facilitate additional network reconfiguration as a function of behavioral requirements.

The lack of behavioral constraints during the resting state make it inherently difficult to directly test whether the predicted alterations in phasic catecholaminergic activity were indeed related to changes in network topology. Fortunately, we could interrogate this hypothesis by leveraging the relationship between the locus coeruleus and the descending sympathetic circuitry that controls pupil dilation^18^, which in turn has been linked to behaviorally relevant alterations in cortical arousal^20–22^. In a previous study, we demonstrated a positive relationship between pupil diameter and fluctuations in network topology^1^, suggesting that ascending neuromodulatory signals may facilitate inter-regional coordination, and hence, network-level integration. In the current study, we hypothesized that the increase in free catecholamines following atomoxetine^23^ should heighten this relationship, and hence lead to a stronger relationship between pupil diameter and network- and regional-level integration. Consistent with this hypothesis, we observed a stronger relationship between pupil diameter and network topology following ATX administration than following placebo (Figure 1c/d). Together, these results provide evidence to suggest that during quiescence, network topology is sensitive to both phasic and tonic patterns of ongoing noradrenergic activity.

### The effect of atomoxetine on the topological signature of cognitive function

A potential benefit of increasing the concentration of free noradrenaline^24^ is that the liberated catecholamines can be utilized in appropriate contexts to facilitate activity within task-relevant neural circuits. In other words, ATX may down-regulate tonic noradrenergic release during rest, but when required, it may conversely facilitate an increase in phasic noradrenergic release^9^ and hence, increase network-level integration. To directly test this hypothesis, we analyzed data from a separate dataset of 19 subjects (age range 18-30) who underwent a cognitively-challenging, parametric N-back task after the administration of either ATX (60mg) or placebo^13^. We hypothesized that, due to a heightened phasic noradrenergic response, the main effect of network-level integration should be more pronounced during the task following ATX, when compared to the placebo condition.

Consistent with our hypothesis, we observed a significant increase in network-level integration during task performance following ATX (Figure 2a). Specifically, there was an inverted U-shaped relationship between cognitive load and network integration in both conditions that was significantly elevated in the post-ATX session (t = 2.47; p = 0.009; Figure 2b). This main effect of ATX was maximal across frontal, parietal and temporal cortices, along with thalamus, amygdala and Crus II of the cerebellum (Figure 2c; red). Importantly, there is a long-standing research literature linking catecholamines with cognitive function via an inverse-U shaped relationship^11,25^, however few studies have provided a potential explanation for the algorithmic benefits that such a mechanism might confer. Here, we demonstrate that network integration may provide one potential mechanism underlying the inverted U-shaped relationship between catecholamine levels and cognitive performance.

**Figure 2.**
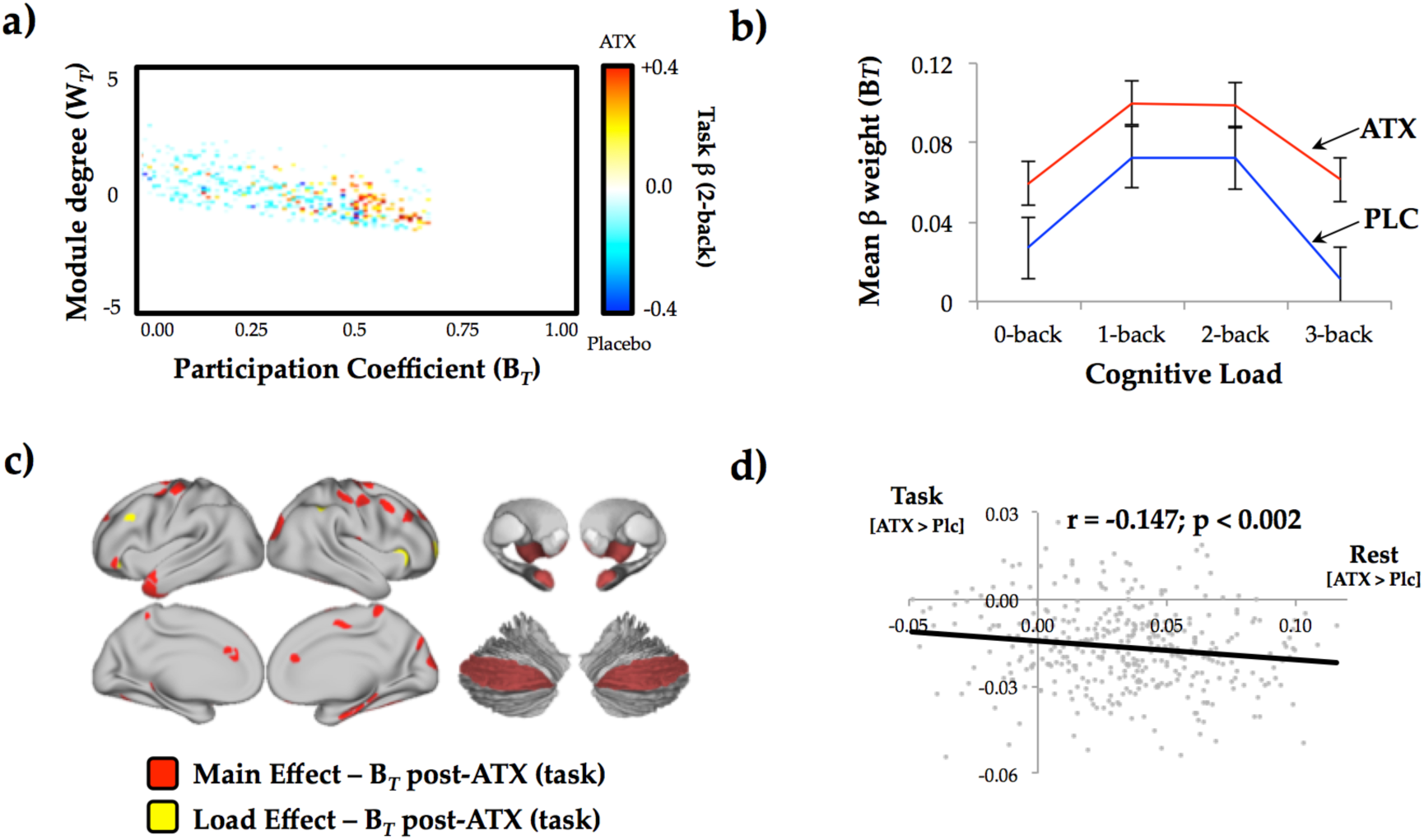
– a) mean cartographic profile across all four blocks of load comparing post-ATX to post-placebo – similar patterns were observed in each block (FDR q ≤ 0.05); b) mean parcel-wise B for each N-back load in both the placebo (PLC; blue) and atomoxetine (ATX; red) conditions (error bars represent standard error across subjects); c) parcels with higher B_*T*_ post-ATX as a function of task performance (FDR q ≤ 0.05) – main effect (red) and load effect (yellow); d) correlation between the regions that showed highest B_*T*_ during task performance (ATX > Placebo) and regions that were shifted towards segregation in the rest study (ATX_(Post>Pre)_ > Placebo_(Post>Pre)_) – see Table S1 for parcel MNI co-ordinates (FDR q ≤ 0.05). Key: ATX – atomoxetine; B_*T*_ – between-module connectivity; W_*T*_ – within-module connectivity.

### Regional topological signatures change as a function of cognitive load

Although the majority of regions across the brain demonstrated an inverse U-shaped relationship with load, there was a subset of regions that demonstrated a linear increase with cognitive load following atomoxetine administration (Figure 2c; yellow). Specifically, the bilateral anterior insula, left dorsolateral prefrontal cortex and right frontopolar cortex demonstrated a higher extent of integration (B_*T*_) with increasing task complexity following ATX, suggesting that the additional free catecholamines may have facilitated enhanced topological involvement of these regions as a function of task performance. Together, the relationship between these regions and ATX suggests that phasic noradrenaline may selectively enhance performance in task-relevant regions, perhaps through arousal-mediated alterations in neural gain^10,26,27^.

### Regional mediation of noradrenergic effects on network topology

The topological dissociation across the two studies analyzed here begs the question – is the effect of ATX on network topology mediated by a set of similar regions across behavioral states? When we directly compared the effect of ATX in the two datasets, we observed a spatial correspondence between the effects of ATX on network topology during rest and task. Specifically, the regional topological signature observed during the N-back task was inversely correlated with the regional signature observed during rest (r = −0.147; p < 0.002; Figure 2d), suggesting that ATX impacted similar regions during rest and task, albeit by shifting them in different topological directions.

## Discussion

Our results provide direct evidence that the manipulation of catecholamine levels in the human brain leads to substantial shifts in network topology. Further, we were also able to demonstrate that the alterations in network topology critically depend on behavioral state. In the resting state, an over-abundance of free catecholamine levels following ATX administration was associated with a relatively segregated network topology (Figure 1a). The lack of effortful cognitive engagement during the resting state may have facilitated a decrease in ascending arousal via ATX-mediated auto-inhibition of the locus coeruleus^6,9^, allowing the network to settle into a segregated architecture, potentially as a way to minimize energy expenditure^28^. In contrast, when presented with a complex behavioral challenge following ATX^13^, an increased phasic-tonic ratio of noradrenergic function^9^ may have facilitated functional connectivity between otherwise segregated circuits, integrating the functional connectome (Figure 2a) and putatively increasing the temporal coordination between the brain circuitry required to successfully complete the N-back task. Together, these results thus provide evidence that the ascending arousal system mediates the balance between network-level integration and segregation as a function of cognitive demands.

The biological mechanism underlying these effects is currently a topic of active investigation, but there is emerging evidence that the network-level impact of catecholamines may relate to their ability to modulate the excitability and gain of neurons across the brain^7,29^. For instance, it has previously been shown that stimulation of the locus coeruleus, both using electrical^30^ and optogenetic approaches^31^, leads to the widespread activation of the cortex, producing a high-frequency, low-amplitude signature that is a known correlate of the awake brain^32^. Similar patterns have also been observed during spontaneous activity in awake mice, confirming that firing in the locus coeruleus directly facilitates high frequency cortical activity during natural behavior^17^. Recent computational modeling work has suggested that the presence of these activated states is crucially dependent on the activation of fast-spiking interneurons in the cortex^33^. Importantly, these interneurons rely on input from ascending neuromodulatory systems, such as the locus coeruleus^34^, to facilitate the synchronous oscillations between activated regions in the gamma frequency^35^, which in turn shape the temporal flow of information processing in the cortex^36^. Together with our findings, these studies further confirm the role of catecholaminergic tone in simultaneously balancing the key topological properties of integration and segregation in a state-dependent manner. They also provide a mechanistic explanation for the brain’s response to periods of acute stress, which are also mediated by ascending noradrenergic systems^19^. Future studies will play an important role in solidifying this mechanistic explanation and determining the contexts in which the balance between these factors is most crucial for understanding complex behavior.

Although our experimental results suggest a crucial role for noradrenaline in the topological reconfiguration of brain network architecture, it bears mention that biological systems rarely demonstrate sharp boundaries between function systems. For instance, in addition to modulating noradrenaline, ATX administration has also been shown to modulate the central concentrations of other arousal-related neurotransmitters, including serotonin^37^, histamine^38^ and dopamine^11^, suggesting that the effects observed in our study may relate to the reconfiguration of the ascending arousal system as a whole. This systemic interdependence is perhaps best exemplified when comparing the relationship between noradrenaline and dopamine, the two major catecholaminergic neurotransmitters in the central nervous system. While the majority of dopaminergic synapses utilize their own specific transporter^39^, a sub-group of dopaminergic terminals in the cortex can also exploit noradrenergic transporters to re-enter pre-synaptic axons^40^. In addition, it has been shown that locus coeruleus neurons can co-release noradrenaline and dopamine^41^. As such, our results may reflect the combined improvements in cortical signal-to-noise that relate to some combination of dopaminergic and noradrenergic effects on neuronal projection targets^11^. The different concentrations of ATX used in the two studies may also have impacted upon these non-selective aspects of ATX. Fortunately, future studies that contrast the roles of the related neurotransmitter systems at different concentrations across a range of behavioral states will help to clarify this issue.

Together, our results demonstrate a relationship between network topology and catecholaminergic function that is sensitive to behavioral state. Future experiments should now be designed to decipher the relative impact of other neurotransmitter systems, both in health and disease.

## Online Methods

### General

All data were taken from two previously-published studies^12,13^. The analysis and hypotheses for the resting state study were pre-registered (https://osf.io/utqq2/), and the code used to analyze the data is freely available at http://github.com/macshine/.

## Resting State Study

### Participants

24 right-handed individuals (age 19 –26 years; 5 male) were included in this study. All participants were screened by a physician for physical health and drug contraindications. Exclusion criteria included: standard contraindications for MRI; current use of psychoactive or cardiovascular medication; a history of psychiatric illness or head trauma; cardiovascular disease; renal failure; hepatic insufficiency; glaucoma; hypertension; drug or alcohol abuse; learning disabilities; poor eyesight; smoking >5 cigarettes a day; and current pregnancy. All participants gave written informed consent before the experiment.

### Study Design

We used a double-blind placebo-controlled crossover design^12^. In each of two sessions, scheduled 1 week apart at the same time of day, participants received either a single oral dose of atomoxetine (40 mg) or placebo (125 mg of lactose monohydrate with 1% magnesium stearate, visually identical to the drug). In both sessions, participants were scanned once before pill ingestion (t = −20 min) and once following ingestion (t = 90 min), when approximate peak-plasma levels are reached. Each scan comprised 8 min of eyes-open resting-state fMRI. During scanning, the room was dark, and participants fixated on a black fixation cross presented on a gray background. Drug uptake was confirmed using cortisol and *α*-amylase levels in the saliva^12,23^.

### MRI Data

All MRI data were collected with a Philips 3T MRI scanner. In each of the scanning sessions, we collected T2*-weighted EPI resting-state images (echo time 30 ms, repetition time 2.2 s, flip angle 80°, FOV 80 x 80 x 38 voxels of size 2.75 mm isotropic, and 216 volumes). To allow magnetic equilibrium to be reached, the first 5 volumes were automatically discarded. In addition, each time the participant entered the scanner, we collected a B0 field inhomogeneity scan (echo time 3.2 ms, repetition time 200 ms, flip angle 30°, and FOV 256 x 256 x 80 voxels with a reconstructed size of 0.86 x 0.86 mm with 3-mm-thick slices). Finally, at the start of the first session, we collected a high-resolution anatomical T1 image (echo time 4.6 ms, repetition time 9.77 ms, flip angle 8°, and FOV 256 x 256 x 140 voxels with size 0.88 x 0.88 mm with 1.2-mm-thick slices) for image normalization and registration.

### Data Preprocessing

After realignment (using FSL’s MCFLIRT) and skull stripping (using BET), B0 unwarping was used to control for potential differences in head position across sessions. The B0 scans were first reconstructed into an unwrapped phase angle and magnitude image. The phase image was then converted to units of radians per second and median filtered, and the magnitude image was skull-stripped. We then used FEAT to unwarp the EPI images in the y-direction with a 10% signal loss threshold and an effective echo spacing of 0.333. The un-warped EPI images were then pre-whitened, smoothed at 5 mm FWHM, and co-registered with the anatomical T1 to 2 mm isotropic MNI space (degrees of freedom: EPI to T1, 3; T1/EPI to MNI, 12). FMRIB’s ICA-based X-noiseifier^44^ was used with pre-trained weights to denoise the imaging data.

Temporal artifacts were identified in each dataset by calculating framewise displacement (FD) from the derivatives of the six rigid-body realignment parameters estimated during standard volume realignment^45^, as well as the root mean square change in BOLD signal from volume to volume (DVARS). Frames associated with FD > 0.25mm or DVARS > 2.5% were identified, however as no participants were identified with greater than 10% of the resting time points exceeding these values, no trials were excluded from further analysis. There were no differences in head motion parameters between the four sessions (p > 0.500). Following artifact detection, nuisance covariates associated with the 12 linear head movement parameters (and their temporal derivatives), DVARS, physiological regressors (created using the RETROICOR method) and anatomical masks from the CSF and deep cerebral WM were regressed from the data using the CompCor strategy^46^. Finally, in keeping with previous time-resolved connectivity experiments^47^, a temporal band pass filter (0.071 < f < 0.125 Hz) was applied to the data.

### Brain Parcellation

Following pre-processing, the mean time series was extracted from 375 pre-defined regions-of-interest (ROI). To ensure whole-brain coverage, we extracted: 333 cortical parcels (161 and 162 regions from the left and right hemispheres, respectively) using the Gordon atlas^48^, 14 subcortical regions from Harvard-Oxford subcortical atlas (bilateral thalamus, caudate, putamen, ventral striatum, globus pallidus, amygdala and hippocampus; http://fsl.fmrib.ox.ac.uk/), and 28 cerebellar regions from the SUIT atlas^49^ for each participant in the study.

### Time-Resolved Functional Connectivity

To estimate functional connectivity between the 375 ROIs, we used the Multiplication of Temporal Derivatives approach (MTD; http://github.com/macshine/coupling/^14^). The MTD is computed by calculating the point-wise product of temporal derivative of pairwise time series (equation 1). In order to reduce the contamination of high-frequency noise in the time-resolved connectivity data, the MTD is averaged by calculating the mean value over a temporal window, *w*. Time-resolved functional connectivity was calculated between all 375 brain regions using the MTD within a sliding temporal window of 15 time points (~33 seconds), which allowed for estimates of signals amplified at approximately 0.1 Hz. Individual functional connectivity matrices were then calculated within each temporal window, thus generating an un-thresholded (that is, signed and weighted) 3-dimensional adjacency matrix (region × region × time) for each participant.

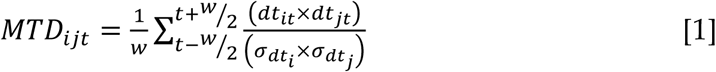

**Equation 1** – Multiplication of Temporal Derivatives, where for each time point, *t*, the MTD for the pairwise interaction between region *i* and *j* is defined according to equation 1, where *dt* is the first temporal derivative of the *i*^th^ or *j*^th^ time series at time *t*, σ is the standard deviation of the temporal derivative time series for region *i* or *j* and *w* is the window length of the simple moving average. This equation can then be calculated over the course of a time series to obtain an estimate of time-resolved connectivity between pairs of regions.

### Time-Resolved Community Structure

The Louvain modularity algorithm from the Brain Connectivity Toolbox (BCT^15^ was used in combination with the MTD to estimate both time-averaged and time-resolved community structure. The Louvain algorithm iteratively maximizes the modularity statistic, *Q*, for different community assignments until the maximum possible score of *Q* has been obtained (equation 2). The modularity estimate for a given network is therefore a quantification of the extent to which the network may be subdivided into communities with stronger within-module than between-module connections.

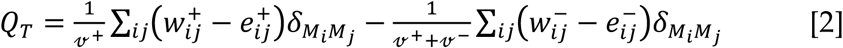

**Equation 2** – Louvain modularity algorithm, where *v* is the total weight of the network (sum of all negative and positive connections), *w_ij_* is the weighted and signed connection between regions *i* and *j*, *e_ij_* is the strength of a connection divided by the total weight of the network, and *δ_MiMj_* is set to 1 when regions are in the same community and 0 otherwise. ‘+’ and ‘−’ superscripts denote all positive and negative connections, respectively.

For each temporal window, the community assignment for each region was assessed 500 times and a consensus partition was identified using a fine-tuning algorithm from the Brain Connectivity Toolbox (http://www.brain-connectivity-toolbox.net/). This then afforded an estimate of both the time resolved modularity (Q_*T*_) and cluster assignment (Ci_*T*_) within each temporal window for each participant in the study. To define an appropriate value for the γ parameter, we iterated the Louvain algorithm across a range of values (0.5 – 2.5 in steps of 0.1) for 100 iterations of a single subjects’ time-averaged connectivity matrix and then estimated the similarity of the resultant partitions using mutual information. A γ parameter of 1.1 provided the most robust estimates of topology across these iterations.

### Cartographic Profiling

Based on time-resolved community assignments, we estimated within-module connectivity by calculating the time-resolved module-degree Z-score (W_*T*_; within module strength) for each region in our analysis (equation 3^50^).

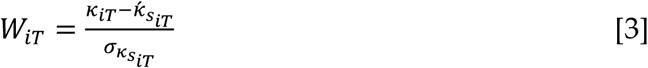

**Equation 3** – Module degree Z-score, W*_iT_*, where κiT is the strength of the connections of region *i* to other regions in its module s_i_ at time *T*, *κ́_s_iT__* is the average of κ over all the regions in s*i* at time *T*, and *σ_κ_s_iT___* is the standard deviation of κ in si at time *T*.

To estimate between-module connectivity (B*_T_*), we used the participation coefficient, B*_T_*, which quantifies the extent to which a region connects across all modules (i.e. between-module strength; equation 4).

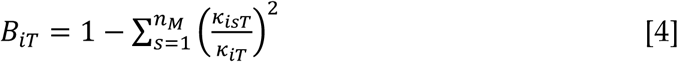

**Equation 4** – Participation coefficient B*_iT_*, where κ_isT_ is the strength of the positive connections of region *i* to regions in module *s* at time *T*, and κ_iT_ is the sum of strengths of all positive connections of region *i* at time *T*. The participation coefficient of a region is therefore close to 1 if its connections are uniformly distributed among all the modules and 0 if all of its links are within its own module.

To track fluctuations in cartography over time, for each temporal window, we computed a joint histogram of within- and between-module connectivity measures, which we refer to here as a ‘cartographic profile’ (Figure 1a). Code for this analysis is freely available at https://github.com/macshine/integration/.

### Pupilometry

Pupil size was measured from the right eye at 500 Hz with an MRI-compatible Eyelink 1000 eye tracker. Blinks and other artifacts were interpolated offline using shape-preserving piecewise cubic interpolation. Pupil data were low-pass filtered at 5 Hz to remove high-frequency noise and Z-scored across conditions. Five participants were excluded from pupil-related analyses due to poor signal quality (≥50% of continuous time series interpolated) or missing data. Of the remaining participants, on average 20% ± 9% of the data were interpolated. We replicated our previous observation^1^ of a positive relationship between pupil diameter and integrated network topology (Figure S1a).

### Null Model Creation

To determine whether the integrative signature of the brain was more dynamic than predicted by a stationary null model (Figure S1b^51^, surrogate data was created using a stationary Vector Auto Regressive model (order was set at 6 to match the expected temporal signature of the BOLD response in 2.2s TR data). The mean covariance matrix across the entire experiment was used to generate 2,500 independent null data sets, which allows for the appropriate estimation of the tails of non-parametric distributions^52^. These time series were then preprocessed using the same approach outlined for the BOLD data. For each analysis, we estimated the kurtosis of the mean B_*T*_ time series for each of the 2,500 simulations. We then calculated the 95^th^ percentile of this distribution and used this value to determine whether the resting state data fluctuated more frequently than the null model. We found that the dynamic network structure within the fMRI data had a higher kurtosis than the 95^th^ percentile of the stationary null model (Figure S1b), confirming the presence of non-stationarity.

### Statistical Analyses

The following hypotheses were pre-registered with the Open Science Framework (https://osf.io/utqq2/):

#### Hypothesis 1

To explicitly test whether the resting brain fluctuates more frequently than a stationary null model, we will calculate the kurtosis of the window-to-window difference in the mean BT score for each iteration of a vector autoregression (VAR) null model (model order = 6). The mean covariance matrix across all 24 subjects from the pre-placebo session will be used to generate 2,500 independent null datasets, which allows for the appropriate estimation of the tails of non-parametric distributions^52^. These time series will then be filtered in a similar fashion to the BOLD data. For each analysis, we will create a statistic for each independent simulation that summarized the extent of fluctuations in the null dataset. We will then calculate the 95th percentile of this distribution and used this value to determine whether the resting state data fluctuated more frequently than the null model (i.e. whether there deviations as extreme as the 95th percentile of the null dataset occur more than 5% of the time).

#### Hypothesis 2

We will estimate the Spearman’s rho correlation between the convolved pupil diameter and the time series of each bin of the cartographic profile. We will then fit a linear mixed-effects model with random intercept to determine whether the correlation between each bin of the cartographic profile is more extreme than chance levels (FDR q ≤ 0.05).

#### Hypothesis 3

Group level differences will be investigated by comparing each bin of the cartographic profile for all subjects prior to and post-atomoxetine administration, as well as prior to and post-placebo using a 2x2 ANOVA design. Specifically, the percentage of time that each bin of the cartographic profile is occupied during the resting state for each of the four sessions will be entered into a 2x2 ANOVA. Our hypothesis predicts a significant interaction effect between pre- and post-placebo and pre- and post-atomoxetine administration. We will correct for multiple comparisons using a false discovery rate of q ≤ 0.05. Similar interaction effects will be assessed at the regional level (i.e. regional topological measures, such as participation coefficient) using a series of 2x2 ANOVA designs.

### Task-based Study

Based on our interim results, we hypothesized that, if phasic and tonic noradrenaline release differentially alter the balance between integration and segregation, then integration should be stronger following ATX during cognitive task performance. This hypothesis was not identified as part of our pre-registration, but arose as a *post hoc* interrogation of the data. To test this hypothesis, we analyzed data from a different dataset, in which 19 participants (age: 18-30; all right-handed males) underwent a cognitively-challenging N-back task following either ATX (60mg; a higher dose than utilized in the first study) or placebo in a double-blind, randomized placebo-controlled crossover design (PLC-ATX n=8; ATX-PLC n=11^13^). The study was carried out in accordance with the Declaration of Helsinki and was approved by the local medical ethics committee of Maastricht University Medical Centre (NL53913.068.15). All participants gave written informed consent prior to each session and were reimbursed for participation.

### Behavioral Task

Participants performed a parametrically modulated N-back task, in which they were required to identify target letters that were presented for 1000ms on an LCD screen during MRI scanning. Targets consisted of letters that were the same as the letter presented one, two or three trials previously (i.e. 1-back, 2-back, or 3-back, respectively). A further control condition was also involved, in which participants were asked to detect the letter ‘X’ (i.e. 0-back). Every task condition was presented three times in pseudo-random order (3-4 targets per block). Participants responded to targets and distractors with right index and middle finger button presses, respectively. Task effects were modeled for fMRI analysis using a block design.

### MRI data and preprocessing

All MRI data were collected with a Siemens 3T MRI scanner. In each of the scanning sessions, we collected T2*-weighted EPI images (echo time 30 ms, repetition time 2.0 s, flip angle 77°, and 286 volumes). To allow magnetic equilibrium to be reached, the first 5 volumes were automatically discarded. In addition, each time the participant entered the scanner, we collected a B0 field inhomogeneity scan (echo time 3.2 ms, repetition time 200 ms, flip angle 30°, and FOV 256 x 256 x 80 voxels with a reconstructed size of 0.86 x 0.86 mm with 3-mm-thick slices). Finally, at the start of the first session, we collected a high-resolution anatomical T1 image (echo time 4.6 ms, repetition time 9.77 ms, flip angle 8°, and FOV 256 x 256 x 140 voxels with size 0.88 x 0.88 mm with 1.2-mm-thick slices) for image normalization and registration. Data were preprocessed in a similar fashion to the resting-state analysis, albeit with a more liberal upper bound on the band pass filter (0.01 < *f* < 0.2 Hz), in order to account for potential task-related alterations in functional connectivity, and without correction for physiological parameters, which were not collected in the study.

### fMRI analysis

Preprocessed BOLD data were subjected to the same time-resolved network analysis pipeline as to the one utilized for the resting state analysis. Following this step, both regional (W_*T*_ and B_*T*_) and global (cartographic profile) time series were modeled against the blocks of the 0-, 1-, 2- and 3-back conditions in both the post-ATX and post-placebo sessions. Instruction screens, rest blocks and head motion parameters were also modeled. We then statistically compared the resultant β weights for each of the blocks separately using a series of F-tests (FDR q ≤ 0.05) with the following two contrasts: i) main effects, which were modeled as the mean activity in the 1-, 2- and 3-back blocks versus the 0-back block; and ii) load effects, which were modeled as a parametric increase in activity as a function of cognitive load across the four blocks. None of the effects were significantly correlated with head motion, either within- or between-subjects (p>0.500).

Finally, we correlated the β weights for the main effect of ATX > PLC during the task for each parcel with the interaction effect of ATX_[Post>Pre]_ > PLC_[Post>Pre]_ on resting-state topology using a Pearson’s correlation (the task data did not contain a “pre-drug” condition). The significance of this correlation was determined by randomly permuting the task-based effects 5,000 times and then re-estimating the Pearson’s correlation between the shuffled effects and the original regional effects in the resting state. The inverse correlation between the two parcel values was more extreme than the 0.02^nd^ percentile of the null distribution.

## References

1. Shine, J. M. et al. The Dynamics of Functional Brain Networks: Integrated Network States during Cognitive Task Performance. Neuron 92, 544–554 (2016).

2. Vatansever, D., Menon, D. K., Manktelow, A. E., Sahakian, B. J. & Stamatakis, E. A. Default Mode Dynamics for Global Functional Integration. J. Neurosci. 35, 15254–15262 (2015).

3. Kitzbichler, M. G., Henson, R. N. A., Smith, M. L., Nathan, P. J. & Bullmore, E. T. Cognitive effort drives workspace configuration of human brain functional networks. J. Neurosci. 31, 8259–270 (2011).

4. de Pasquale, F., Penna, Della, S., Sporns, O., Romani, G. L. & Corbetta, M. A Dynamic Core Network and Global Efficiency in the Resting Human Brain. Cereb. Cortex bhv185 (2015). doi:10.1093/cercor/bhv185

5. Samuels, E. R. & Szabadi, E. Functional neuroanatomy of the noradrenergic locus coeruleus: its roles in the regulation of arousal and autonomic function part II: physiological and pharmacological manipulations and pathological alterations of locus coeruleus activity in humans. Curr Neuropharmacol 6, 254–285 (2008).

6. Sara, S. J. The locus coeruleus and noradrenergic modulation of cognition. Nat. Rev. Neurosci. 10, 211–223 (2009).

7. Aston-Jones, G. & Cohen, J. D. An Integrative Theory of Locus Coeruleus-Norepinephrine Function: Adaptive Gain and Optimal Performance. Annu. Rev. Neurosci. 28, 403–450 (2005).

8. Eldar, E., Cohen, J. D. & Niv, Y. The effects of neural gain on attention and learning. Nat Neurosci (2013).

9. Bari, A. & Aston-Jones, G. Atomoxetine modulates spontaneous and sensory-evoked discharge of locus coeruleus noradrenergic neurons. Neuropharmacology 64, 53–64 (2013).

10. Devilbiss, D. M. & Waterhouse, B. D. Phasic and tonic patterns of locus coeruleus output differentially modulate sensory network function in the awake rat. Journal of Neurophysiology 105, 69–87 (2011).

11. Robbins, T. W. & Arnsten, A. F. T. The Neuropsychopharmacology of Fronto-Executive Function: Monoaminergic Modulation. Annu. Rev. Neurosci. 32, 267–287 (2009).

12. van den Brink, R. L. et al. Catecholaminergic Neuromodulation Shapes Intrinsic MRI Functional Connectivity in the Human Brain. J. Neurosci. 36, 7865–7876 (2016).

13. Hernaus, D., Casales Santa, M. M., Offermann, J. S. & Van Amelsvoort, T. Noradrenaline transporter blockade increases fronto-parietal functional connectivity relevant for working memory. Eur Neuropsychopharmacol 27, 399–410 (2017).

14. Shine, J. M. et al. Estimation of dynamic functional connectivity using Multiplication of Temporal Derivatives. NeuroImage 122, 399–407 (2015).

15. Rubinov, M. & Sporns, O. Complex network measures of brain connectivity: Uses and interpretations. NeuroImage 52, 1059–1069 (2010).

16. Mattar, M. G., Cole, M. W., Thompson-Schill, S. L. & Bassett, D. S. A Functional Cartography of Cognitive Systems. PLoS Comput. Biol. 11, e1004533 (2015).

17. Safaai, H., Neves, R., Eschenko, O., Logothetis, N. K. & Panzeri, S. Modeling the effect of locus coeruleus firing on cortical state dynamics and single-trial sensory processing. Proc. Natl. Acad. Sci. U.S.A. 112, 12834–12839 (2015).

18. Nieuwenhuis, S., De Geus, E. J. & Aston-Jones, G. The anatomical and functional relationship between the P3 and autonomic components of the orienting response. Psychophysiology 48, 162–175 (2011).

19. Hermans, E. J., van Marle, H. & Ossewaarde, L. Stress-related noradrenergic activity prompts large-scale neural network reconfiguration. Science 334, 1151–3 (2011).

20. McGinley, M. J., David, S. V. & McCormick, D. A. Cortical Membrane Potential Signature of Optimal States for Sensory Signal Detection. Neuron 87, 179–192 (2015).

21. Joshi, S., Li, Y., Kalwani, R. M. & Gold, J. I. Relationships between Pupil Diameter and Neuronal Activity in the Locus Coeruleus, Colliculi, and Cingulate Cortex. Neuron 89, 221–234 (2016).

22. Reimer, J. et al. Pupil fluctuations track rapid changes in adrenergic and cholinergic activity in cortex. Nat Commun 7, 13289 (2016).

23. Warren, C. M., van den Brink, R. L., Nieuwenhuis, S. & Bosch, J. A. Norepinephrine transporter blocker atomoxetine increases salivary alpha amylase. Psychoneuroendocrinology 78, 233–236 (2017).

24. Invernizzi, R. W. & Garattini, S. Role of presynaptic *α*2-adrenoceptors in antidepressant action: recent findings from microdialysis studies. Progress in Neuro-Psychopharmacology and Biological Psychiatry 28, 819–827 (2004).

25. Cools, R. & D’Esposito, M. Inverted-U-shaped dopamine actions on human working memory and cognitive control. Biological Psychiatry 69, e113–25 (2011).

26. Waterhouse, B. D., Moises, H. C. & Woodward, D. J. Noradrenergic modulation of somatosensory cortical neuronal responses to lontophoretically applied putative neurotransmitters. Experimental Neurology 69, 30–49 (1980).

27. Reimer, J. et al. Pupil Fluctuations Track Fast Switching of Cortical States during Quiet Wakefulness. Neuron 84, 355–362 (2014).

28. Bullmore, E. & Sporns, O. The economy of brain network organization. Nat. Rev. Neurosci. 13, 336–349 (2012).

29. McGinley, M. J. et al. Waking State: Rapid Variations Modulate Neural and Behavioral Responses. Neuron 87, 1143–1161 (2015).

30. Toussay, X., Basu, K., Lacoste, B. & Hamel, E. Locus coeruleus stimulation recruits a broad cortical neuronal network and increases cortical perfusion. J. Neurosci. 33, 3390–3401 (2013).

31. Carter, M. E. et al. Tuning arousal with optogenetic modulation of locus coeruleus neurons. Nat Neurosci 13, 1526–1533 (2010).

32. Berridge, C. W. & Waterhouse, B. D. The locus coeruleus–noradrenergic system: modulation of behavioral state and state-dependent cognitive processes. Brain Research Reviews 42, 33–84 (2003).

33. Stringer, C. et al. Inhibitory control of correlated intrinsic variability in cortical networks. Elife 5, 91 (2016).

34. Castro-Alamancos, M. A. & Gulati, T. Neuromodulators produce distinct activated states in neocortex. J. Neurosci. 34, 12353–12367 (2014).

35. Cardin, J. A. et al. Driving fast-spiking cells induces gamma rhythm and controls sensory responses. Nature 459, 663–667 (2009).

36. Constantinidis, C., Williams, G. V. & Goldman-Rakic, P. S. A role for inhibition in shaping the temporal flow of information in prefrontal cortex. Nat Neurosci 5, 175–180 (2002).

37. Ding, Y. S. et al. Clinical doses of atomoxetine significantly occupy both norepinephrine and serotonin transports: Implications on treatment of depression and ADHD. NeuroImage 86, 164–171 (2014).

38. Liu, L. L., Yang, J., Lei, G. F. & Wang, G. J. Atomoxetine Increases Histamine Release and Improves Learning Deficits in an Animal Model of Attention-Deficit Hyperactivity Disorder. Basic & Clinical Pharmacology and Toxicology 102, 527–32 (2008).

39. Chen, N. & Reith, M. Structure and function of the dopamine transporter. European journal of pharmacology 405, 329–39 (2000).

40. Morón, J. A., Brockington, A. & Wise, R. A. Dopamine uptake through the norepinephrine transporter in brain regions with low levels of the dopamine transporter: evidence from knock-out mouse lines. Journal of Neuroscience 22, 389–95 (2002).

41. Devoto, P., Flore, G., Pira, L. & Longu, G. Alpha2-adrenoceptor mediated co-release of dopamine and noradrenaline from noradrenergic neurons in the cerebral cortex. Journal of Neuroscience 88, 1003–9 (2004).

42. Carbonell, F., Nagano-Saito, A., Leyton, M. & Cisek, P. Dopamine precursor depletion impairs structure and efficiency of resting state brain functional networks. Frontiers in Neuroinformatics 84, 90–100 (2014).

43. Achard, S. & Bullmore, E. Efficiency and Cost of Economical Brain Functional Networks. PLoS Comput. Biol. 3, e17 (2007).

44. Salimi-Khorshidi, G., Douaud, G. & Beckmann, C. F. Automatic denoising of functional MRI data: combining independent component analysis and hierarchical fusion of classifiers. NeuroImage 90, 449–68 (2014).

45. Power, J. D. et al. Methods to detect, characterize, and remove motion artifact in resting state fMRI. NeuroImage 84, 320–341 (2014).

46. Behzadi, Y., Restom, K., Liau, J. & Liu, T. T. A component based noise correction method (CompCor) for BOLD and perfusion based fMRI. NeuroImage 37, 90–101 (2007).

47. Bassett, D. S., Yang, M., Wymbs, N. F. & Grafton, S. T. Learning-induced autonomy of sensorimotor systems. Nat Neurosci 18, 744–751 (2015).

48. Gordon, E. M. et al. Generation and Evaluation of a Cortical Area Parcellation from Resting-State Correlations. Cereb. Cortex 26, 288–303 (2014).

49. Diedrichsen, J., Balsters, J. H., Flavell, J., Cussans, E. & Ramnani, N. A probabilistic MR atlas of the human cerebellum. NeuroImage 46, 39–46 (2009).

50. Guimerà, R. & Nunes Amaral, L. A. Functional cartography of complex metabolic networks. Nature 433, 895–900 (2005).

51. Laumann, T. O., Snyder, A. Z., Mitra, A. & Gordon, E. M. On the stability of bold fmri correlations. Cerebral Cortex EPub Ahead of Print (2016).

52. Nichols, T. E. & Holmes, A. P. Nonparametric permutation tests for functional neuroimaging: a primer with examples. Hum Brain Mapp 15, 1–25 (2002).

